# Resolution of structural variation in diverse mouse genomes reveals chromatin remodeling due to transposable elements

**DOI:** 10.1101/2022.09.26.509577

**Authors:** Ardian Ferraj, Peter A. Audano, Parithi Balachandran, Anne Czechanski, Jacob I. Flores, Alexander A. Radecki, Varun Mosur, David S. Gordon, Isha A. Walawalkar, Evan E. Eichler, Laura G. Reinholdt, Christine R. Beck

**Author notes:** **Corresponding Author and Lead Contact:** Christine R. Beck. **Author Contributions** C.R.B. conceived and supervised the study. A.F. performed data preprocessing, data analysis, and data interpretation. P.A.A. performed quality control, method development, and data analysis. A.C. and L.G.R. performed cell culture work, sample preparation, and sample submission for sequencing. P.B. and I.A.W. aided in sample submission, data management, and preprocessing. J.I.F., A.A.R., and V.M. designed primers and conducted PCRs for validation work. D.S.G. and E.E.E. generated and provided the GRCm39 segmental duplication track. A. F., P.A.A., and C.R.B. wrote the manuscript.

## Abstract

Diverse inbred mouse strains are among the foremost models for biomedical research, yet genome characterization of many strains has been fundamentally lacking in comparison to human genomics research. In particular, the discovery and cataloging of structural variants is incomplete, limiting the discovery of potentially causative alleles for phenotypic variation across individuals. Here, we utilized long-read sequencing to resolve genome-wide structural variants (SVs, variants ≥ 50 bp) in 20 genetically distinct inbred mice. We report 413,758 site-specific SVs that affect 13% (356 Mbp) of the current mouse reference assembly, including 510 previously unannotated variants which alter coding sequences. We find that 39% of SVs are attributed to transposable element (TE) variation accounting for 75% of bases altered by SV. We then utilized this callset to investigate the impact of TE heterogeneity on mouse embryonic stem cells (mESCs), and find multiple TE classes that influence chromatin accessibility across loci. We also identify strain-specific transcription start sites originating in polymorphic TEs that modify gene expression. Our work provides the first long-read based analysis of mouse SVs and illustrates that previously unresolved TEs underlie epigenetic and transcriptome differences in mESCs.

## INTRODUCTION

Mice of varying genetic backgrounds are often utilized to create models of human disease and to generate populations of infinitely diverse substrains that exhibit a wide range of genotypic and phenotypic heterogeneity. Such panels include the Collaborative Cross (CC) and Diversity Outbred (DO) populations, which are multiparental groups of recombinant inbred lines and derivative outbred stocks constructed from eight original founder strains (Churchill et al., 2004; Threadgill et al., 2011). These reference panels are often used to determine genotype-phenotype relationships and have been invaluable tools for precise genomic mapping of numerous quantitative trait loci, including regions associated with addiction, stem cell pluripotency, and insulin secretion (Bubier et al., 2014; Keller et al., 2019; Skelly et al., 2020).

Discovery of the genetic origins of numerous diseases and phenotypic characteristics present within diverse populations depends on high-quality reference genomes and precise variant catalogs (Nurk et al., 2022). For decades now, genetic and genomic studies have depended on the *mus musculus* reference genome, which is derived from the popular C57BL/6J strain. Although the C57BL/6J (GRCm39) reference is the most complete mouse assembly to date, reliance on an assembly built from one genetic background limits the analysis of genetic variation among diverse populations as sequencing reads from divergent haplotypes are often misplaced or unmapped. This problem has been noted in human research (Audano et al., 2019), but is compounded in mouse, where there is a much larger amount of genetic diversity in laboratory inbred strains than there is in the human population. Despite recent efforts to construct strain-specific reference genomes (Lilue et al., 2018), much of the genomic landscape that is variable between these strains remains incomplete due to the technological limitations of short-read whole genome sequencing. Regions that remain unresolved include complex repeats, segmental duplication-mediated variants, and transposable element variants, which are often represented as discontinuous sequences in current strain-specific reference builds. These regions can contain important sequences or variants that affect genes and potentially lead to phenotypic change. Until we can accurately resolve them, the underlying impact of these structural variants (SVs) on mouse biology remains unknown.

Recent advances in long-read sequencing have allowed for accurate resolution of repetitive DNA and large insertion variants (Chaisson et al., 2019; Ebert et al., 2021; Nurk et al., 2022) and have greatly increased sensitivity for SVs over short-read technologies. In particular, detailed studies of the same genomes in humans have revealed that 60-70% of SVs are missed when relying solely on short-read sequence data, and for those detected by both platforms, only the long-read sequence platform can fully resolve the allele (Chaisson *et al*., 2019; Ebert *et al*., 2021). Although large-scale studies utilizing long-read sequencing in distinct human populations have been published (Ebert *et al*., 2021), to date there have been no efforts to fully resolve and detect SVs across diverse mouse genomes with these technologies.

Here, we utilize Pacific Biosciences (PacBio) long-read whole-genome sequencing to assemble the genomes of 20 diverse inbred laboratory strains of mice. From whole-genome comparisons, we have generated a sequence-resolved callset of 413,758 SVs (54% novel) spanning three subspecies of *mus musculus* (*domesticus, musculus*, and *castaneus*) and up to ∼0.5M years of divergence, including SVs in regions which cannot be resolved with short-read sequencing. We used this resource to identify and resolve transposable element polymorphisms present between diverse mice, revealing multiple retrotransposition competent subfamilies, such as the L1MdTf and IAP types, that cause widespread changes in mouse embryonic stem cell (mESC) chromatin accessibility and gene expression. We present these data as a comprehensive mouse SV resource that can be used for future genomic studies, aid in modeling and studying the effects of genetic variation, and enhance genotype-to-phenotype research.

## RESULTS

### Long-read sequencing assemblies of diverse mouse genomes

Whole-genome long-read PacBio sequencing data was generated for 20 diverse inbred mouse strains to a minimum of 30-fold coverage. We selected a mixture of classical and wild-derived inbred laboratory substrains including the parental founders of the Collaborative Cross (CC) (129S1/SvImJ, A/J, CAST/EiJ, NOD/ShiLtJ, NZO/HILtJ, PWK/PhJ, and WSB/EiJ), six resultant CC animals which harbor extreme phenotypic abnormalities of unknown genetic origin (CC005, CC015, CC032, CC055, CC060, CC074) (Kolmogorov et al., 2019), and six additional substrains with distinct genetic backgrounds (BALB/cByJ, BALB/cJ, C3H/HeJ, C3H/HeOuJ, C57BL/6NJ, PWD/PhJ). We chose this cohort as it represents a wide variety of strains with diverse genetic backgrounds, strains with complex phenotypes of unknown cause, and strains interesting for mapping variants responsible for quantitative traits (Chesler et al., 2008; Gan et al., 2020; Threadgill *et al*., 2011).

We generated *de novo* assemblies for each genome reaching a 6.92 Mbp contig N50 (Table S1) and on average achieve a 534×increase in contig N50 compared to previous short-read assemblies built from the same strains (14 kbp) (Lilue *et al*., 2018). Additionally, our assemblies are contained in 143×fewer contigs (average of 2,483 per genome vs 355,353) with 228 Mbp of additional sequence on average compared to short-read assemblies. Therefore, we have created the most contiguous genome assemblies of diverse mouse genomes produced to date.

### Structural variants are prevalent across mouse genomes

For each of the 20 strains, we aligned each strain de novo assembly to the GRCm39 (mm39, mus musculus) reference genome and called SVs with the phased assembly variant caller (PAV) (Ebert *et al*., 2021). From these strain-specific SV calls, we created a non-redundant variant callset by merging SVs across strains (Ebert *et al*., 2021); In total we detected 413,758 SVs which occur at unique sites across the current mouse reference assembly (Table S2), composed of 244,859 (59%) insertions, 168,652 (41%) deletions, and 247 (0.06%) inversions (Figure 1A). Strains within the *domesticus* subspecies are more closely related to the C57BL/6J reference strain; we find that these strains contain the fewest variant calls per animal, with an average of 60,490 SVs per mouse genome (Table 1). Strains from the *musculus* (PWD/PhJ, & PWK/PhJ) and *castaneus* (CAST/EiJ) subspecies contain an average of 200,401 SVs per genome when compared to the C57BL/6J reference. We calculate a 5.8% false discovery rate from 584 PCR validations (73 SVs across 8 samples) (Table S3). We identify 97,901 (24%) SVs which are unique to a single substrain and 1,483 shared variants (Figure 1A) that indicate reference biases or rare SVs found within the reference (Audano *et al*., 2019). We also find 268,436 SVs (243 Mbp) specific to one subspecies (Figure 1B), of which 73% (178 Mbp of insertion sequence) are absent from GRCm39, indicating genomic variants that are inaccessible without sequencing technology capable of spanning SVs.

**Figure 1.**
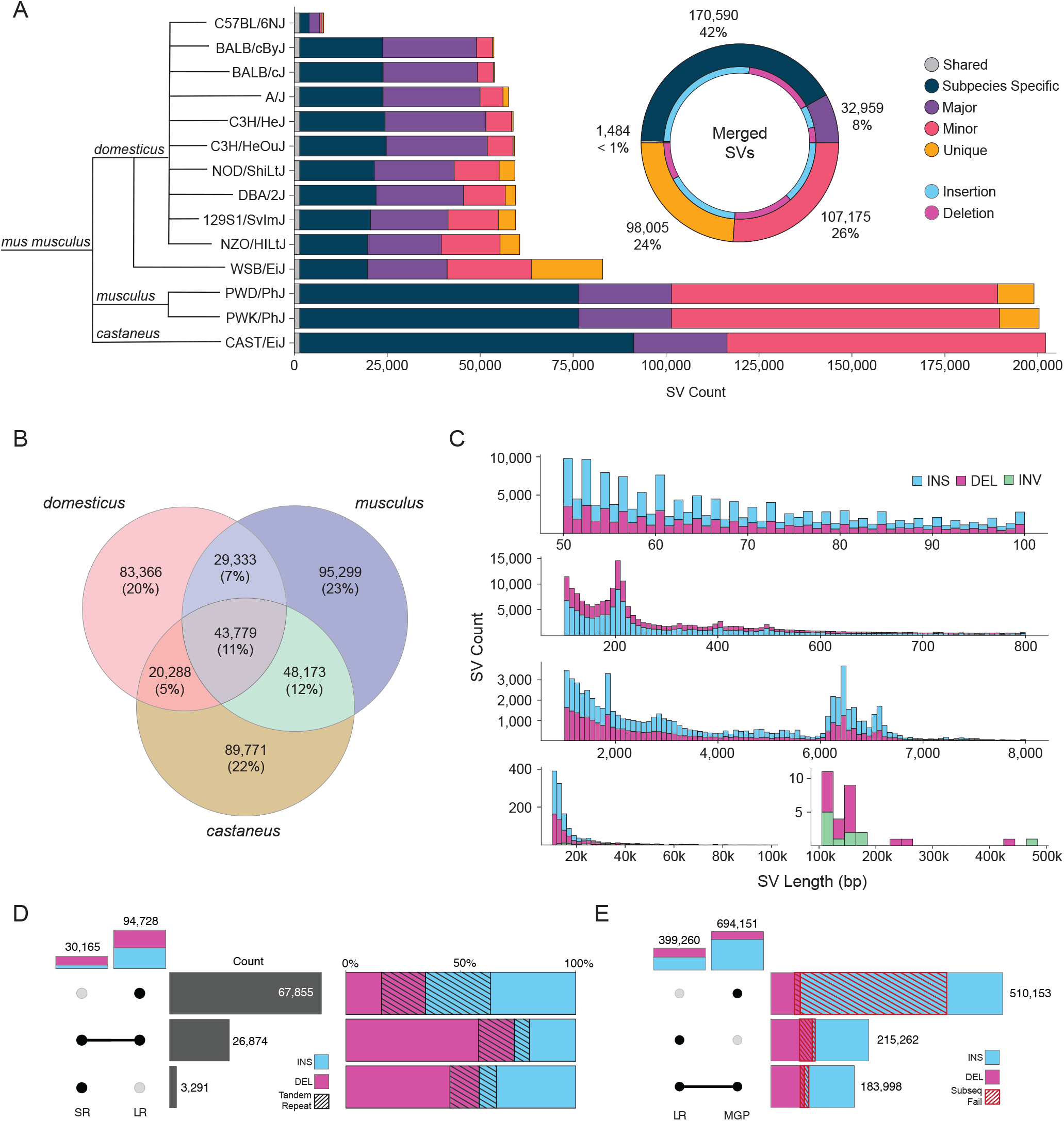
Discovery of Structural Variants in Diverse Mouse Genomes. (A) Total number of structural variants discovered in each mouse genome when compared to the *mus musculus* (GRCm39, C57BL/6J background) reference genome. Variants are grouped by their frequency within the cohort: shared (present in all strains) Major (≥ 50% of the cohort), Minor (< 50%), exclusive to one strain (Unique) or exclusive to one subspecies (Subspecies specific). We merged variants into a non-redundant callset (donut plot), show along with the proportion of insertion and deletion calls (blue and purple). (B) Total number of SVs discovered and shared between each*-* subspecies sequenced (*domesticus, musculus*, and *castaneus*). (C) Length distribution of structural variants identified as insertions (blue), deletions (purple), and deletions (green). (D) Average number of SVs per mouse genome supported by long read (LR) and short read (SR) detection, along with the proportion of insertions and deletions which contain tandem repeat sequences (black striped bar). (E) Total number of SVs from long read (LR) and mouse genomes project (MGP) callset, along with the proportion of variants validated by raw long-read alignment size differences (Red striped bar).

The amount of variation that these SVs contribute to is extensive; the 416,758 SVs encompass a total of 356 Mbp of variable sequence, accounting for 13% of the current mouse reference assembly. We find that these SVs contain 4.9×the number of bases affected when compared to previously published single nucleotide variants from diverse mouse genomes (Keane et al., 2011). The length of SVs varies greatly, and the size of SVs are non-randomly distributed (Figure 1C). An increased number of small SVs (50-100 bp) contain variants with lengths divisible by 2 resulting from dinucleotide repeat expansions and contractions; this has previously been observed in human studies (Ebert *et al*., 2021). We also observe peaks in the size distribution consistent with abundant transposable element polymorphisms (200 bp, SINEs; 6.3 kbp, LINEs). Although the majority of variants are under 1 kbp in length (83%), the total sequence content due to SVs is dominated by those 1 kbp or greater, which account for 292 Mbp (82%) of variable sequence. Each species contains an average of 81 Mbp of unique sequence due to SVs. We found that a large portion of SVs (201,342, 49%) were found within tandemly repeated regions.

When compared to human genomes, which have an average of 24,653 SVs per individual (Ebert *et al*., 2021), we find 60,490 SVs per *domesticus* genome (2.3×human), indicating greater diversity from SVs among mouse genomes than human genomes. Because they are determined by comparing *de novo* assemblies to a *domesticus* reference, the number of SVs per genome is even greater in *musculus* (199,597, 8.1×human) and *castaneus* (202,011, 8.2×human) substrains. With extensive structural polymorphism across mouse genomes, the use of a single linear reference may therefore be inadequate for mapping genomic data, especially from more diverged strains. For example, per genome, we find that SV insertions duplicate whole genes (20 in *domesticus*, 37 in *castaneus*, and 28 in *musculus*), suggesting that the effect of paralogous gene copies has been systematically underestimated by short-read approaches due to reference biases and the limitations of a single linear reference.

### Long-read sequencing data reveals novel SVs

Previously, a number of studies used short-read sequencing data to interrogate SVs across large cohorts of genetically diverse inbred mice (Doran et al., 2016; Keane *et al*., 2011; Lilue *et al*., 2018; Srivastava et al., 2017). To identify the subset of variation from our study that was previously undefined and to add orthogonal support for our assembly-based SV callset, we performed SV calling from Illumina whole-genome sequencing of 18 previously sequenced substrains that overlapped our cohort (Lilue *et al*., 2018; Srivastava *et al*., 2017). When we compare the SV callsets from long-read and short-read data, on average 91% of short read deletion calls and 84% of insertions were detected in our long-read SV callset (Figure 1D). Conversely, short-read SV calling was only able to detect 46% of deletions, 14% of insertions, and 39% of inversions discovered by long-read sequencing. Across the 18 strains, SV calling from long reads detected an additional distinct 213,688 insertions, 64,277 deletions, and 97 inversions. Notably, short-read sequencing is particularly underpowered for the detection of SVs in repeat regions (Chaisson *et al*., 2019; Zhao et al., 2021); the number of SVs which are supported by short reads drops considerably for both deletions (46% to 24%) and insertions (14% to 9%) when considering variants within tandem repeat regions. In total, 155,156 simple repeat SVs were not detected by short reads.

To compare our SV resource with previously published variant calls, we intersected our long-read callset with the most recent mouse genomes project (MGP) SVs (Doran *et al*., 2016; Keane *et al*., 2011). Overall, 54% (215,262) of SVs we call are novel to our study (Figure 1E). Interestingly, intersection of the long-read callset with the MGP dataset revealed there were a large number of insertions (445,538) which were not detected by long-read methods (Figure 1E). To determine the validity of these MGP specific calls, we mapped long reads generated from each sample to the GRCm39 reference genome and surveyed each SV region for changes in sequence length (mean read length difference ≥ 50 bp). From this we determined that 95% (117,715) of the insertions specific to long-read detection were supported by raw long-read alignments, while only 28% (122,981) of the MGP-only insertions were supported. This leaves a total of 322,574 (72%) potentially false insertion SV calls within the MGP SV dataset, while only 6,372 (5%) of PAV-only insertions were not supported by raw read alignments. Calls detected only with long-reads were significantly smaller (mean of 425 bp vs 1393 bp for MGP-supported calls (*p < 0*.*0001*, Student’s *t*-test). This was also true for variants which contain simple repeat sequences (mean of 279 bp for novel variants vs 1631 bp for MGP-supported calls; *p < 0*.*0001*, Student’s *t*-test). These data suggest that short-read variant calling falters in the detection of smaller SVs.

### Long-read assemblies reveal extensive transposon variation at a nucleotide level

The most notable improvement from using long-read genome assemblies comes from the ability to reconstruct long repetitive regions. This is particularly imperative when characterizing mouse genetic variation as mice contain elevated retrotransposon insertion rates when compared to human (Feusier et al., 2019; Richardson et al., 2017). In order to create a more complete resource and investigate the impact of mobile element variation in diverse mice, we utilized our SV callset to identify transposable element variants (TEVs) between each mouse genome. We find that 39% (162,787) of SVs between all samples are attributable to TEVs, with most of the TEVs being insertions (97,100, 60%). TEVs are dominated by LINE-1 polymorphisms (47%, 76,640), followed by SINEs (B1 & B2; 24%, 39,389), and various endogenous retroviruses (ERVs) (ERVK, ERVL-MaLR, ERVL, ERV1; 28%, 45,204) (Figure 2A). We observe various peaks of TEV length, consistent with the variant calls representing retrotransposition events. We observe various modes of TEV length consistent with retrotransposition, with accumulations of TEVs at ∼200 bp (SINEs), ∼7.2 kbp (ERVK LTRs) and a ∼6.3 kbp peak for full length LINE-1s (Figure 2B). In addition to classifying major TE families, we further annotate TEVs by subtype; We find that *mus musculus* LINE-1 variants are predominantly of the L1MdA (26,717 variants) subtype, followed by L1MdTf (17,010), and L1MdGf (10,649), fitting previous studies which have detailed active LINE-1 subfamilies in *mus musculus* (Jachowicz and Torres-Padilla, 2016). In addition to comprising 39% of nonredundant variant sites, TEV s constitute 75% (278 Mbp) of the sequence content structurally variable among mouse genomes (Figure 2C). LINE-1 variants alone constitute 47% (172 Mbp) of the varied sequence content, in contrast with 24% (92 Mbp) of variable sequence contributed by non-TEVs. Although SINEs comprise 25% of TE variant sites, they only account for 2.1% (7.9 Mbp) of variable sequence.

**Figure 2.**
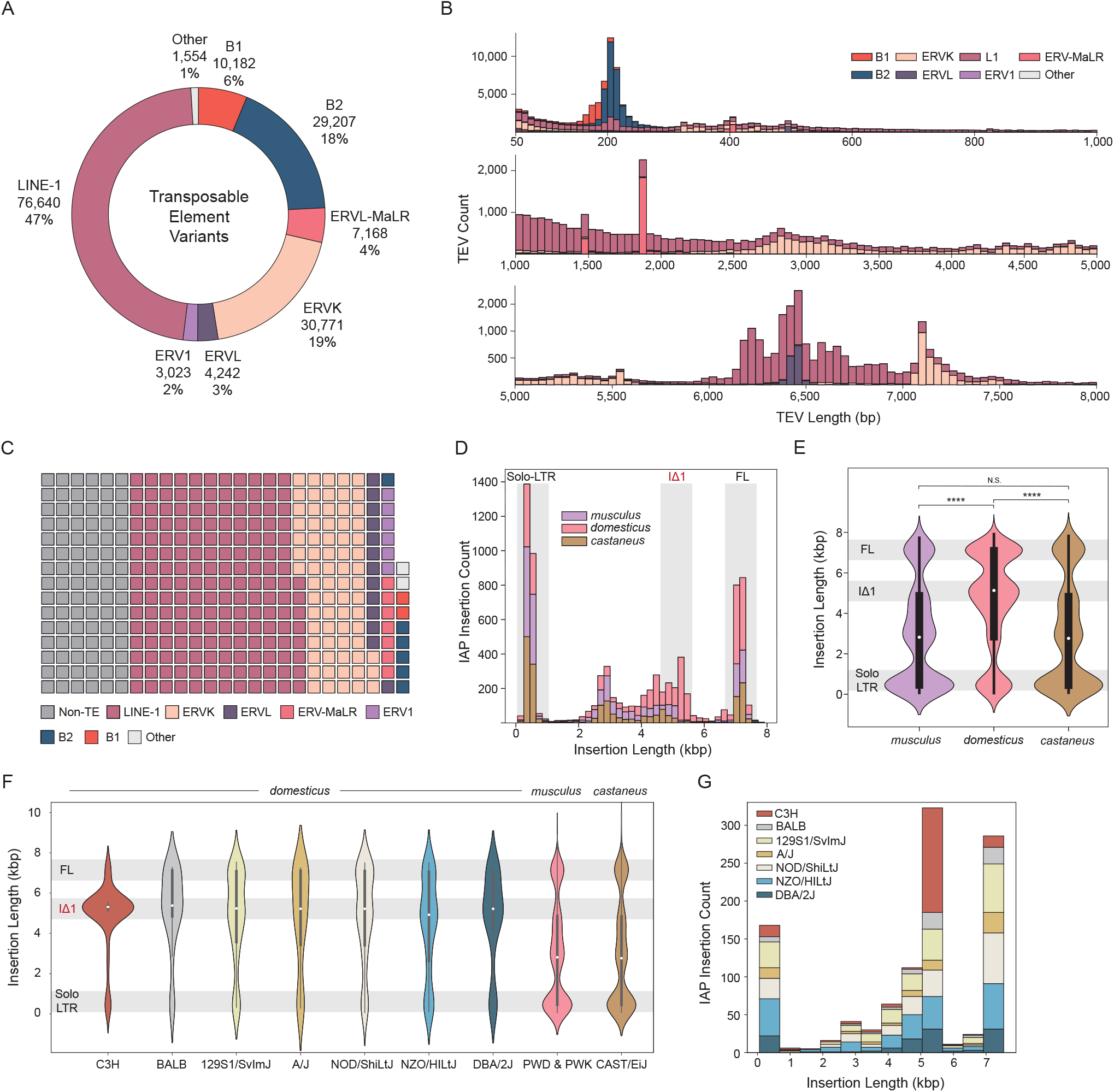
Transposable Element Variation in Diverse Mouse Genomes. (A) Total number of transposable element variants (TEVs) discovered from long-read sequencing by major transposon family. (B) Length distribution of TEVs by major transposon family. (C) Total number of variable bases due to TEV by major transposon families and non-TE structural variants. Each block equals one megabase of variable sequence. (D) Count of subspecies-specific intracisternal A particle (IAP) variants. Highlighted size ranges are solo-LTR (∼300 bp), IΔ1 variants (4.5-5.5 kbp), and full length (FL) variants (6.5-7.5 kbp). (E) Size distribution of species-specific IAP insertions (Mann Whitney U test; ****: p ≤ 1e-4). (F) Size distribution of strain-specific IAP insertions within the domesticus lineage. Substrains which share a parental origin (C3H: C3H/HeJ & C3H/HeOuJ, BALB: BALB/cByJ & BALB/cJ) are grouped to represent each lineage. (G) Count of strain-specific IAP insertions in closely related domesticus animals.

When compared to all SVs, we observe, as expected, fewer TEVs within and around genic regions (43% of intergenic variants are TEVs vs 26% non-intergenic, Table S4). Coding sequence and intronic variants are also underrepresented for TEVs. At the repeat family level, we find LINE-1 variants to be enriched in intergenic regions and depleted for coding sequence, intronic, UTR, and nearby gene regions. Conversely, SINE elements (mainly tRNA derived B2s) tend to accumulate in and around genic regions, with significant enrichment in introns, UTRs and up and downstream regions. This enrichment was not observed with the 7SL derived SINE B1 elements. Various ERVs are depleted in introns; however, splice donor site variants are significantly enriched for ERVL and ERV1 sequences when compared to other SVs. We find that 41% of all ERV variants that contain splice donor sites are from the MT retrotransposon family, which are known to contribute to chimeric RNA sequences and are developmentally regulated (Peaston et al., 2004).

### IAP endogenous retroviral elements have variable rates of insertion in mouse genomes

Intracisternal A-particle (IAP) elements are murine specific retroviral elements which are a known driver of variation in mice (Rebollo et al., 2020). Full length (∼7.2 kbp) IAPs are autonomous long terminal repeat (LTR) retrotransposons which can cause aberrant splicing and disease if they insert near or within genes (Duhl et al., 1994; Morgan et al., 1999). Proper annotation of long terminal repeat (LTR) elements is especially difficult with short reads as many subtypes are differentiated by their internal structure. We utilized our TEV callset to investigate polymorphic IAP LTRs within the *mus musculus* lineage. We find that substrains of the *domesticus* lineage contain 43% (3363 insertions) of all subspecies-specific IAP polymorphisms (Figure 2D). Strains from the *domesticus* subspecies contains a significant increase (*p* ≤ 2.41e-2, Fisher’s exact) in the proportion of IAPs which constitute all ERVK insertions (54%) when compared to *castaneus* (44%) and *musculus* (43%), which is consistent with previous findings that suggest an ongoing and *domesticus*-specific IAP expansion (Nellaker et al., 2012). We further detail this expansion by providing all variable IAP sequences between these subspecies and observe a *domesticus-*specific increase in the proportion of variable bases due to IAP insertions in comparison to other ERVKs (67% from active IAP subtypes, total of 15 Mbp), representing an increase in the variable sequence content of *domesticus* when compared to *castaneus* (56%, 6 Mbp) and *musculus* (56%, 7 Mbp) IAP sequences. IAP insertions in *domesticus* also contain a different nucleotide length distribution, with notable accumulation of full-length elements as well as variants that are within a 4.5-5.5 kbp size range (Figure 2E). These shorter, or intermediate sized sequences are indicative of IΔ1 IAPs, which contain an internal deletion relative to full length elements and remain active in *mus musculus* (Ishihara et al., 2004; Kuff and Lueders, 1988). Here, we show evidence that the expansion of IAP TEs in *domesticus* is driven by both full length and variable length IAPs.

Interestingly, even between closely related *domesticus* substrains we observe a difference in the distribution of IAP lengths. Mice with a parental C3H background contain a notable size discrepancy for IAP insertions which are 4.5-5.5 kbp in length when compared to other *domesticus* strains (Figure 2F). Previous studies have detailed a hypermutable IAP phenotype within the C3H/HeJ substrain, with increased expression of IAPs relative to other mice of closely related background (Rebollo *et al*., 2020). We find that strain C3H/HeJ contains a significant increase in the proportion of variants within the IΔ1 size range when compared to every non-C3H substrain in our study (*p* < 0.05, Fisher’s exact). Within closely related *domesticus* substrains, we find that C3H mice contain over one third of strain-specific IAP insertions within the IΔ1 size range (Figure 2G). Hyperactive IAP expression and numerous gene altering polymorphisms have been well documented in C3H mice (Gagnier et al., 2019; Rebollo *et al*., 2020). Here, we uncover a large number of unique IAP insertions specific to C3H mice and catalog a higher number of insertion polymorphisms when compared to other *domesticus* lines (34% of strain specific insertions in the IΔ1 size range).

### The consequences of structural variation on mouse genomes

To assess the potential functional impact of mouse SV, we utilized the Ensembl Variant Effect Predictor (VEP) tool to intersect SV calls with known genomic features and predict variant severity (McLaren et al., 2016). We find that 55% (228,232) of SVs map within or around (5 kbp up/downstream) genes, with 43% (179,239) intergenic, and 13% (52,411) intersecting mouse regulatory regions (Figure 3A). Most genic SVs (98%) are intronic, with 2% overlapping a non-intron feature (3’ UTR, 5’ UTR, and coding regions) (Figure 3B). We report 829 coding sequence (CDS) variants (512 deletions and 317 insertions), 2,647 3’ UTR variants (1,148 deletions and 1,499 insertions) and 451 5’ UTR SVs (280 deletions and 171 insertions). We then selected all SVs with potential functional consequences and performed an association analysis based on multiple criteria (Figure 3C). First, we determined the tendency of a particular SV class to be detected by short-read sequencing when compared to long reads. Among all CDS variants, 577 (70%) are uniquely called with long reads and 252 (30%) are detected by both long and short reads. We then compared all variant consequences to previously published mouse SVs (Doran *et al*., 2016; Keane *et al*., 2011). Of all variants that were novel to long-read detection, we report 94,863 to be intronic, 1,469 UTR, and 510 coding sequence variants. Interestingly, across all SV calls, coding sequence variants were enriched for simple repeat sequences (Table S4). We additionally conducted 144 PCR validations (18 PCR reactions across eight strains) for SVs which alter coding regions across the eight CC founder strains yielding orthogonal data for SVs leading to frameshift and insertion variants (Table S4).

**Figure 3.**
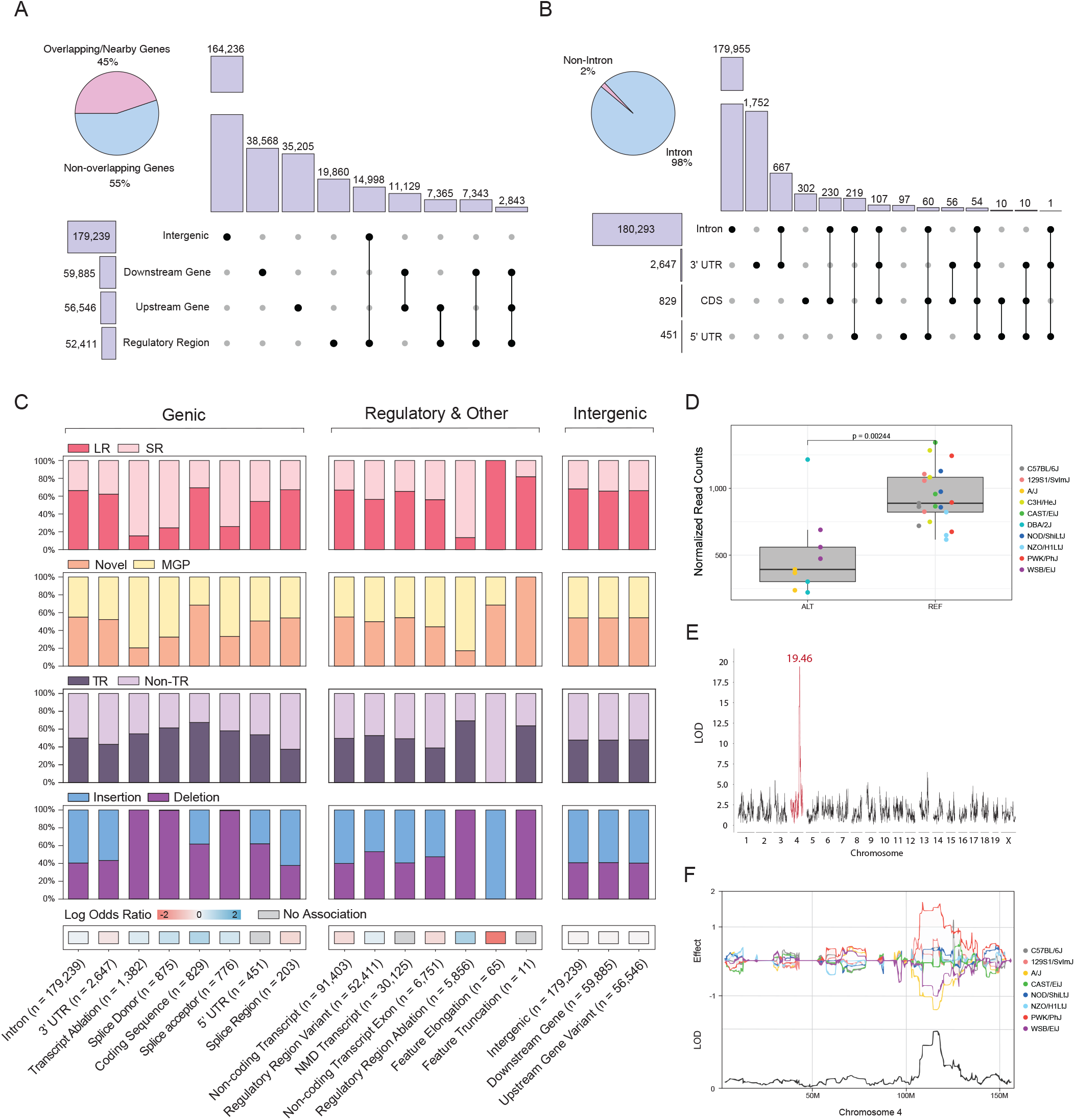
Structural Variant Consequences. (A) Count of structural variants (SVs) overlapping various intergenic regions, with the percentage of variants which overlap genic / non-genic regions. (B) Count of SVs which overlap various genic features, with percentage of variants which overlap a non-intronic or intronic region of a gene. (C) Genic, regulatory, and intergenic SV consequences. Each SV consequence is shown with the proportion that is specific to long read detection (LR: Long Read Only or SR: Short read support), novel to our study when compared to the mouse genomes project database (Novel or MGP), comprised of tandem repeats (Tandem Repeat & Non-Tandem Repeat), and comprised of insertion or deletion SVs (Insertion & Deletion). Each consequence was determined to be correlated or not with a simple repeat SVs (red & blue). (D) *Mutyh* expression differences between strains which contain a 5.3kbp IAP insertion within intron 8 of *Mutyh*. (E) LOD score for a significant cis-eQTL for the *Mutyh* gene. (F) Effect score of each collaborative cross founder strain for the *Mutyh* eQTL on chromosome 4.

Recent studies have detailed a *domesticus*-specific C>A transversion mutator allele associated with a haplotype of the *Mutyh* gene (Sasani et al., 2022), however a causal variant has not been identified. To investigate this allele for potentially causative SVs that are *in cis* with the SNVs on this haplotype, we surveyed each mouse genome for large variants in the *Mutyh* gene locus. We uncovered a 5.3 kbp IAP insertion SV within intron 1 of *Mutyh* that is unique to long-read detection. Furthermore, when we manually examined the haplotypes with the IAP insertion using IGV, the insertion was always on a haplotype that contained the SNVs of the D-like *Mutyh* mutator allele. IAPs have been shown to alter gene structure and negatively impact gene expression through chromatin silencing (Duhl *et al*., 1994; Morgan *et al*., 1999; Sadic et al., 2015). In order to investigate *Mutyh* expression differences of these strains, we performed bulk RNA sequencing on mouse embryonic stem cells (mESCs) derived from 10 substrains (129S1/SvImJ, A/J, C3H/HeJ, CAST/EiJ, DBA/2J, NOD/ShiLtJ, NZO/HILtJ, PWK/PhJ, and WSB/EiJ) and grouped each sample by their insertion status to perform differential gene expression analysis of *Mutyh*. Mice which contained the transposon insertion also contained a significant decrease in *Mutyh* expression compared to those with no IAP insertion (Figure 3D). Interestingly, upon manual curation, all strains that contain the IAP insertion (DBA/2J, A/J, and WSB/EiJ) also contain the five distinct single nucleotide variants previously associated with the mutator phenotype, otherwise known as the “D-like allele” (Sasani *et al*., 2022). We additionally searched previously published mESC eQTL data derived from a large, outbred stock of mice (Diversity Outbred, DO) constructed from the CC founder substrains. We found an extremely significant (LOD score of 19.46) cis-eQTL for the *Mutyh* gene (Skelly *et al*., 2020) (Figure 3E). From this population, samples which contain a negative effect (A/J and WSB/EiJ) match the expression correlation we find with respect to the IAP insertion (Figure 3F). These data show that important regions of the mouse genome can be altered by SVs that are missed in previous studies and show that previously unresolved SVs can reside in import genes and be associated with phenotypic variation.

### Polymorphic transposons alter mESC chromatin dynamics and gene expression

TEs often promote genome diversity and give rise to species-specific neofunctionalization events of biological pathways (Judd et al., 2021; Modzelewski et al., 2021). Furthermore, TE sequences are active in early mouse and human development, and aid in specifying tissue-specific biology (Miao et al., 2020). They inherently harbor promoters and regulatory sequences, therefore we hypothesized that polymorphic TEs would alter the cis regulatory landscape of early development, resulting in altered chromatin accessibility and transcript variation. To investigate the functional impact of these polymorphisms, we utilized our long-read callset to determine the genome-wide impact of TEV on mouse embryonic stem cell chromatin accessibility. We performed ATAC-seq on mESCs derived from 10 genetically diverse substrains (129S1/SvImJ, A/J, C57BL/6J CAST/EiJ, NOD/ShiLtJ, NZO/HILtJ, PWK/PhJ, WSB/EiJ, C3H/HeJ, and DBA/2J) to detect open chromatin regions and surveyed small (5 kbp) regions surrounding SVs in each mouse to detect variant associations with chromatin accessibility (Figure 4A). From this, we found 22,123 (14%) TEVs which are associated with a significant change (*p* < 0.05, MannWhitney Wilcoxon Test) in chromatin accessibility.

**Figure 4.**
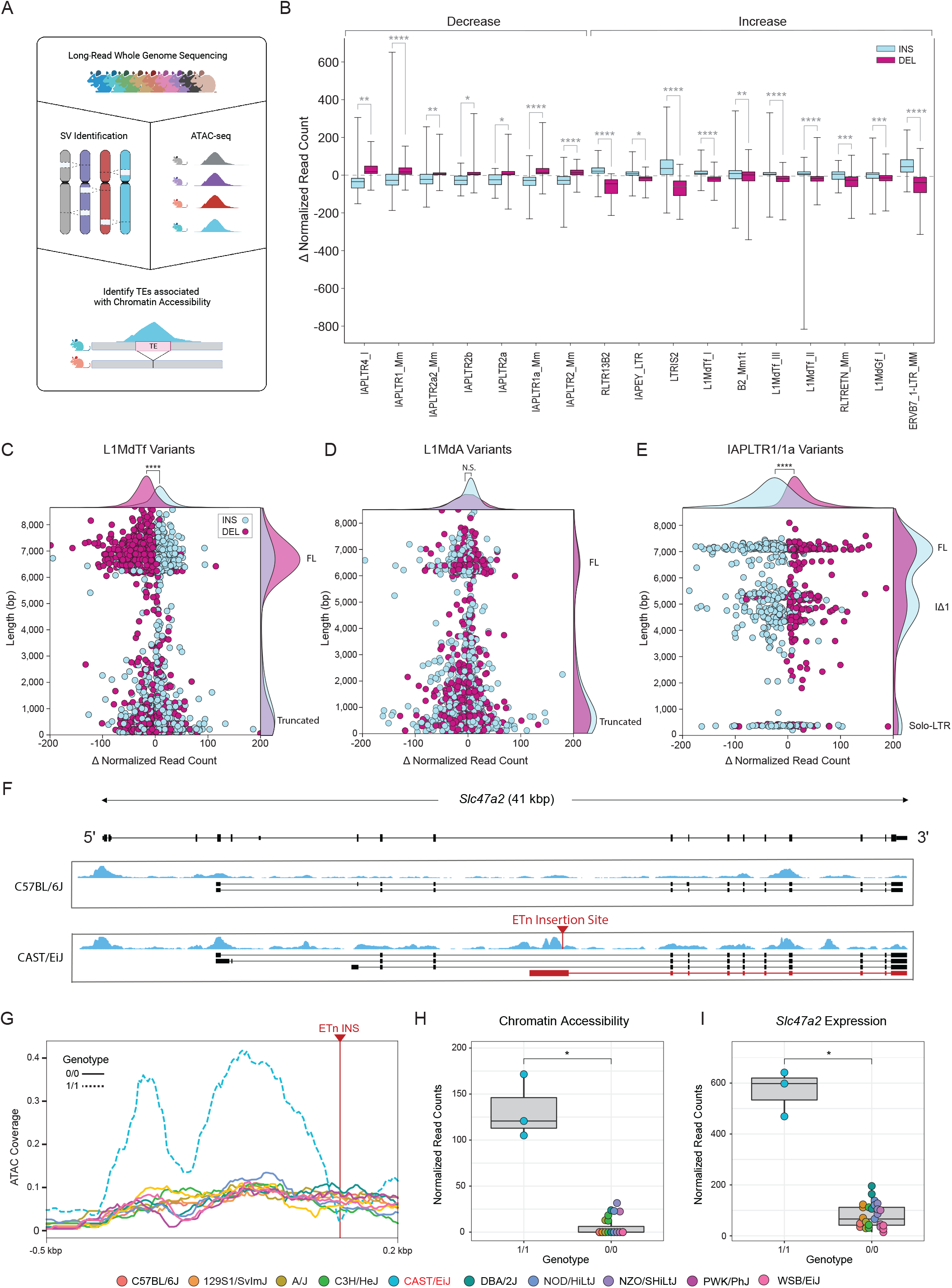
Transposable Element Variant Effects on Chromatin Accessibility and Gene Expression. (A) Diagram showing experimental approach; Long-read whole genome sequencing of diverse mouse genomes allow for comprehensive identification of transposable element variation (TEV). Open chromatin regions found from ATAC-sequencing was used to profile changes in chromatin accessibility at sites of TEVs. (B) Change in chromatin accessibility (normalized ATAC coverage) for 17 transposable element subtypes whose insertion status (insertion: blue, deletion: purple) correlated with changes in chromatin accessibility (Mann Whitney U test; *: 1e-2 < p ≤ 5e-2; **: 1e-3 < p ≤ 1e-2; ***: 1e-4 < p ≤ 1e-3; ****: p ≤ 1e-4). (C) Change in chromatin accessibility (normalized ATAC coverage) plotted against TEV length for L1MdTf variants. Each point represents a L1MdTf variant, categorized by insertion (blue) and deletion (purple). Two distinct clusters of length represent full length (FL) and truncated elements. (D) Change in chromatin accessibility (normalized ATAC coverage) plotted against TEV length for L1MdTf variants. (E) Change in chromatin accessibility (normalized ATAC coverage) plotted against TEV length for IAPLTR1/1a variants. Three distinct clusters of length represent full length (FL), IΔ1, and solo-LTR elements. (F) Genome browser diagram showing gene *Slc47a2*. Strain-specific mESC RNA transcriptome assemblies for strains C57BL/6J and CAST/EiJ are shown with ATAC-Seq signal (blue). Site of a *de novo* ETn insertion specific to strain CAST/EiJ is shown (red arrow) and is accompanied by an alternative transcript not found in C57BL/6J (red transcript). (G) ATAC coverage at the site of CAST/EiJ ETn insertion (H) mESC ATAC-Seq coverage profiles of ten strains surrounding the ETn insertion site between CAST/EiJ and all other strains which do not contain the insertion. (I) Changes in *Slc47a2* gene expression between CAST/EiJ and all other strains which do not contain the insertion.

To further investigate if specific TEVs are responsible for altered chromatin accessibility, we grouped TEVs by family and subtype to determine if the insertion status of a given element is associated with a change in chromatin accessibility. We identified 17 distinct TE subtypes where there was a significant chromatin accessibility difference between insertions and deletions (Figure 4B) with the most frequent genome-wide changes facilitated by LINE-1 variants. Full-length (∼6.3 kbp) polymorphic L1MdTf elements were associated with strain-specific changes in chromatin accessibility with a cluster of elements that are ∼6 kbp long (Figure 4C), suggesting that the 5’ UTR of L1MdTf elements contains regulatory sequences which promote the formation of euchromatin in mESCs. This distribution was not seen in L1MdA elements and is most likely due to the variation in the 5’ UTR monomeric repeat (Figure 4D). Overall, 18% (2,522 of 16,304) of L1MdTf variants found from the 10 strains were associated with a significant change in chromatin accessibility, compared to 8% (2,048 of 23,863) of L1MdA variants. IAP variants had the opposite effect, with insertions promoting heterochromatin formation (Figure 4E). Three distinct clusters of IAP variants based on size were associated with chromatin variation; however, only two clusters (∼7.2 kbp & ∼5.3 kbp) showed a strong trend in accessibility changes based on insertion status. The third cluster consists of short (∼300 bp) sequences, which are remnants of recombination between the 5’ and 3’ LTR ends. This pattern reflects the effects of the internal structure of the IAP element, which contains a short heterochromatin inducing sequence (SHIN) known to be active in mESCs (Sadic *et al*., 2015). We now show that these sequences are often propagated in the mouse genome, leading to genome-wide changes in chromatin accessibility.

To investigate potential transcriptomic changes due to polymorphic TEs, we utilized our RNA-seq dataset generated from the same cell lines to quantify mESC gene expression relative to nearby SVs and constructed strain specific transcript assemblies. In multiple cases, we observe changes in gene expression and neighboring chromatin accessibility correlated with TEVs. We highlight a strain-specific (CAST/EiJ) ETn (Early Transposon) insertion within intron 9 of the gene *Scl47a2* (Figure 4F). This insertion is accompanied by a differential ATACseq signal unique to CAST/EiJ that flanks the insertion site (Figures 4G and 4H) and elevated levels of *Slc47a2* expression in CAST/EiJ in relation to strains absent of the insertion (Figure 4I). Transcript assemblies revealed a CAST/EiJ specific transcript which initiates at the site of the polymorphic ETn insertion (Figure 4F, red arrow and transcript). These data suggest that the polymorphic ETn insertion introduced a novel alternative transcriptional start site within the canonical intron 9 of *Slc47a2*. ETn sequences are known to contain binding sites for the pluripotency factor *Oct4*; we find that polymorphic ETn sequences are strongly associated with chromatin changes (*p* = 7.72e-21, Students *t*-test) and are often associated with changes in nearby transcription in mESCs.

## DISCUSSION

The laboratory mouse is an important model for mammalian genetics and dissecting the relationship of genotype to phenotype (Griffin et al., 2021; Saul et al., 2019). Genetic reference populations such as the Collaborative Cross and Diversity Outbred mice are reproducible resources of genomic complexity that have led to numerous discoveries of disease-relevant loci, and accurate detection of genetic variation is imperative for finding the causative alleles at these loci. Furthermore, diverse subspecies of *mus musculus* contain an overall greater nucleotide divergence than that of human populations, offering unique advantages to population research (Fujiwara et al., 2022; Schmidt, 2018). However, genetic variation between diverged mice has not been captured in its entirety, stemming from the technological limitations of short-read DNA sequencing, which results in poor sequence discovery and sequence annotation for SVs. SVs are often underrepresented in variant catalogs due to their complexity, association with repetitive regions of genomes, and lack of standardized detection methods (Balachandran and Beck, 2020; Carvalho and Lupski, 2016). Recent advances in long-read whole-genome sequencing have since surpassed short reads in their ability to accurately detect SVs (Zhao *et al*., 2021), and prior to this study, the use of such technology has not been applied to detect SV in diverse mouse genomes. The full potential of mouse genetic reference populations cannot be fully realized until SVs are entirely sequence resolved. We have made important steps in rectifying this deficit.

We characterized and sequence-resolved genome wide SVs present in 20 laboratory mouse strains with longread sequencing technology. This cohort represents various popular research models such as: (1) the parental founders of the CC and DO crosses, which are powerful selective breeding panels used for trait mapping (Iraqi et al., 2008); (2) six resultant CC strains which have interesting phenotypes; (3) a strain often crossed with C57BL/6J that is used for studying genotype-phenotype interactions (Geisert and Williams, 2020; Martins et al., 2021; Sasani *et al*., 2022); and (4) numerous other models which contain unique phenotypes due to genetic background, such as the PWD/PhJ strain, a mouse historically utilized to model hybrid sterility (Chubb and Nolan, 1987; Lustyk et al., 2019). The resulting SV calls can be used for identifying variants associated with disease and to study large segments of DNA which were hidden from previous genomic studies. Here, we find 221 Mbp of insertion sequence not present *mus musculus* reference. These sequences are important for the creation of genetically engineered mice on backgrounds other than C57BL/6J, yet targeted mutagenesis constructs are often constructed using the *mus musculus* reference. For example, substrains from the 129 background are regularly used for genetic engineering, including CRISPR-mediated modification (Qin et al., 2016; Simpson et al., 1997). Efforts to create strain specific assemblies have aimed to correct these biases (Lilue *et al*., 2018), however current assemblies of diverse mouse genomes lack large inserted sequences, duplications, and full annotation of TEs (Ebert *et al*., 2021; Zhao *et al*., 2021). From human studies, we have found that long reads excel in identifying SVs and can be used to create highly contiguous genome assemblies which can also be used for SV detection (Audano *et al*., 2019; Chaisson *et al*., 2019; Ebert *et al*., 2021; Heller and Vingron, 2020). Our study creates strain-specific genome assemblies with greater continuity and comprehensive SV callsets. In total we discovered 278,062 variants which were undetected with short-read SV calling methods, while also capturing 90% of the variants from short-read calls (Figure 1D). We call 215,262 SVs which are absent from public mouse SV catalogs (Doran *et al*., 2016; Yalcin et al., 2011), including 510 coding sequence SVs, indicating the importance of using long-read data (Srivastava *et al*., 2017). Newly emerging tools and techniques can further utilize our callset to genotype large breeding panels, such as the CC, DO, and BXD crosses (Ebler et al., 2022).

An ongoing difficulty in genome assembly is the reconstruction of repetitive element sequences, including TEs. TEV detection in mice is imperative as they contain multiple active families of TEs including ERV, LINE-1, and SINE elements. In addition, mice have a higher *de novo* TE insertion rate than that of humans (∼1 LINE-1 insertion in every 8 live births contrast to 1 in 20-200 live births in humans) (Feusier *et al*., 2019; Richardson *et al*., 2017) and contain retrotransposition competent ERVs (Deniz et al., 2019), therefore the role of TEs in genomic change is an important contributor to murine genomic variation. We find that TEs drive extensive SVs in mice and are the dominant source of variable sequence content in mouse genomes (Figure 2C). By resolving TEVs, we detect subspecies-specific enrichment of variable length IAP accumulation in *domesticus* mice when compared with *castaneus* and *musculus*. Our analysis found IΔ1 and full-length IAP variants significantly contribute to the expansion (Figures 2D and 2E). Previous studies have suggested that there is variation in posttranscriptional processing of IAP-containing sequences between *castaneus* and *domesticus* mice that is due to mutations in the nuclear RNA export factor 1 (Nxf1) protein (Concepcion et al., 2015). We observe that the level of IAP polymorphism in *musculus* substrains is similar what we see in *castaneus* and that both are lower than in *domesticus*, suggesting that variable suppression of IAP elements may have different mechanisms in different subspecies. Even within the *domesticus* lineage, IAP insertions unique to C3H mice contain a distinct length distribution, with a greater number of variants in the 4.5-5.5 kbp range (Figures 2F and 2G). Increased expression of IAP retrotransposons has been singularly observed in mESCs derived from C3H/HeJ mice, with multiple accounts of aberrant gene expression from insertion polymorphisms (Gagnier *et al*., 2019; Rebollo *et al*., 2020). Here we sequence-resolve 187 IAP insertions specific to C3H mice which can be used to further investigate this hyperactive insertion phenotype. Furthermore, fixed transposon sequences can be differentially methylated in the mouse genome and are regulated by trans factors such as KRAB zinc finger proteins (Bertozzi et al., 2020). Fixed TEs can also accumulate mutations which are otherwise blind to short-read detection methods; the longread *de novo* assemblies we provide can be used to investigate allelic and epigenetic heterogeneity within a given repeat and allows for precise mapping of orthogonal sequencing data in a strain-specific manner (Lutz et al., 2003; Salvador-Palomeque et al., 2019).

The biological effects of TEVs are not confined to structural alteration of genomic DNA (Chiang et al., 2017). TEVs frequently carry with them potent regulatory sequences which alter gene expression and chromatin accessibility, sometimes in a tissue-specific manner (Maksakova and Mager, 2005; Maksakova et al., 2009; Miao *et al*., 2020). Furthermore, genomic TEs are increasingly recognized as important contributors to early development as they can be co-opted as enhancers, regions of transcription factor binding sites (Todd et al., 2019), and early expression of TEs can be used to profile early embryonic stages such as zygotic genome activation (Kigami et al., 2003). Our TEV callset and assemblies will enable the examination of the effects of TE polymorphisms on early embryonic tissues and can act as a model for studies in other organisms, including diverse humans. Detailed analysis on genomic and transcriptomic usage of TEs in early development have been performed (Miao *et al*., 2020), however these efforts have focused on the role of fixed TEs present in the *mus musculus* reference. Our study examines polymorphic members of multiple families of TEs, shows that they affect nearby chromatin accessibility, and that they may differentiate mouse transcriptomes in early development. Interestingly, our data suggest that recently transposed events from closely related LINE-1 subfamilies likely differ in the *cis* regulatory elements they bind to. *Mus musculus* LINE-1 subtypes are known to contain alternative promoters constructed from monomeric repeats and are differentially methylated in male germ cell development (Zhou and Smith, 2019). Here, we show that polymorphic L1MdTf elements are more often associated with increases in mESC chromatin accessibility when compared to L1MdA variants despite comprising a lower proportion of total polymorphisms, and this effect is more marked with full length elements, suggesting that the promoter is active in mESCs (Figures 4C and 4D). In contrast, we find that IAPs are associated with chromatin closing, consistent with previous studies which detail an internal short heterochromatin inducing sequence that is active in mESCs (Sadic *et al*., 2015). It’s important to note that we detail these changes at one developmental timepoint, and that TE activity is dynamic during mammalian development (He et al., 2021). Further utilization of this callset with orthogonal data to support additional tissue types will aid in characterizing the effects of TE polymorphisms throughout development.

Several important technological advances have become available in recent years such as ultra-long ONT and PacBio HiFi (Wenger et al., 2019). which have made dramatic improvements in assembly contiguity and assembly-based variant detection (Ebert *et al*., 2021; Nurk *et al*., 2022; Nurk et al., 2020; Wang et al., 2021). While telomere-to-telomere assembly is not yet routinely achievable, complete sequence resolution of diverse mouse genomes may be possible in near future. These technologies will enable phasing for the approximately 5% of mouse genomes that are heterozygous after inbreeding and for wild-derived individuals, and it would more completely resolve segmental duplication loci. Complete assemblies may also enable deeper structural exploration such as genome-wide synteny mapping over 500,000 years of evolution and the examination of rearrangements between complex segmental duplications. The data we present here are an important step towards modernizing mouse genetics, generating a sequence-resolved SV resource, a mESC expression resource, and chromatin accessibility data, which will enable evolutionary research and phenotype-genotype correlations in mice.

## METHODS

### Sample selection

We assembled the genomes of 20 diverse inbred laboratory mouse strains (129S1/SvImJ, A/J, CAST/EiJ, NOD/ShiLtJ, NZO/HILtJ, PWK/PhJ, WSB/EiJ, CC005, CC015, CC032, CC055, CC060, CC074, BALB/cByJ, BALB/cJ, C3H/HeJ, C3H/HeOuJ, DBA/2J, C57BL/6NJ, and PWD/PhJ) with whole-genome long-read sequencing and called SVs in each genome generated in this study. We chose to sequence the founders of the collaborative cross as they represent various diverse genetic backgrounds of *mus musculus* and are utilized to create numerous backcrossed generations of offspring for complex phenotype mapping. We also sequenced six resultant substrains from inbreeding of these founders. Lastly, we sequenced seven additional diverse mice which all carry popular disease phenotypes and additional diversity. Because we do not have orthogonal sequencing data to support genome phasing such as strand seq, our assemblies and SV calls represent a single haplotype. Approximately 95% of inbred mouse genomes are homozygous, and we expect to lose 50% of SVs in residual heterozygous loci. For short-read sequencing analysis, we utilized previously published datasets of Illumina whole-genome sequencing reads (Lilue *et al*., 2018; Srivastava *et al*., 2017).

### mESCs

mESC cell lines from ten diverse mice (129S1/SvImJ, A/J, C57BL/6J CAST/EiJ, NOD/ShiLtJ, NZO/HILtJ, PWK/PhJ, WSB/EiJ, C3H/HeJ, and DBA/2J) were derived and cultured using previously published methods (Czechanski et al., 2014). Cells were thawed onto gelatin in ESM/FBS/2i media and grown to 60-80% confluency. All cell lines were between P11-P13 at the time of the harvest. Cells were dissociated with 0.05% trypsin-EDTA, resuspended in PBS for counting, with 1×10^6^ cells reserved for RNA and 1×10^5^ cells reserved for ATAC-seq. For the RNA sample, the cells were spun down, the supernatant removed, and the pellet was flash frozen and then placed on dry ice. The samples were stored at -80°C until harvest. For the ATAC samples, the cells were spun down and resuspended in 1 ml of freeze media (80% ESM/10% FBS/10%DMSO). The volume was divided in ½ among 2 cryovials. The ATAC samples were placed in Cool Cells in the -80°C and then transferred to LN2 24-48 hours later.

### PacBio long-read whole-genome sequencing

PacBio whole-genome sequencing: Kidney tissue from female mice were supplied from The Jackson Laboratory mouse services (Star Methods). gDNA was first extracted using the Gentra Puregene (Qiagen) kit. Frozen kidney tissues from each mouse were first pulverized using a mortar and pestle and transferred to a 15mL tube containing Qiagen Cell Lysis Solution. Lysate was then incubated with Proteinase K for 3 hours at 55°C, followed by the addition RNase A and continued incubation for 40 minutes at 37°C. Samples were cooled on ice and Protein Precipitation Solution was added. Samples were then vortexed and centrifuged. The supernatant was transferred to a new tube containing isopropanol for precipitation. The remaining pellet was washed with 70% ethanol, air dried, and rehydrated in PacBio Elution Buffer until dissolved. Sample preparation was performed following the continuous long read (CLR) protocol from PacBio using the PacBio SMRTbell Express Template Prep Kit 2.0. 15 µg of high molecular weight gDNA was sheared using 26G Needles for 10-20 passes. Sample showing a broad distribution of DNA 20-100 kbp on Femto Pulse (Agilent) was selected to proceed. Additional shearing was performed to achieve the targeted distribution if needed. Sheared DNA Sample was then concentrated with AMPure PB Beads (PacBio). 10μg of the sheared DNA was end repaired and A-tailed to remove single-strand overhangs and repair DNA damage. This is followed by a ligation to an overhang V3 adapter and clean up with 0.45x AMPure PB. The purified library was subjected to size-selection (> 10kbp) using the Blue Pippin system (Sage Science). The library was purified with 1×AMPure PB library and sequenced on a PacBio Sequel II. Each sample was sequenced to a minimum of 30-fold coverage.

### ATAC sequencing (ATAC-seq)

ATAC-seq (Buenrostro et al., 2015) data was generated from mESCs derived from 10 mouse strains (129S1/SvImJ, A/J, C57BL/6J CAST/EiJ, NOD/ShiLtJ, NZO/HILtJ, PWK/PhJ, WSB/EiJ, C3H/HeJ, and DBA/2J). ATAC-seq libraries were prepared using 50,000 cells as previously described (Corces et al., 2016) with the following modifications: pelleted cells were washed in 150 μl PBS before the addition transposase mixture; transposition reactions were agitated at 1000 rpm; transposed DNA was purified using a Genomic DNA Clean & Concentrator (Zymo); purified DNA was eluted in 21 µl elution buffer (10 mM Tris-HCl, pH 8); transposed fragments were amplified using 2×KAPA HiFi HotStart ReadyMix (Roche) and Nextera DNA CD Indexes (Illumina) for 10 cycles of PCR; PCR reactions were purified using 1.7×KAPAPure beads (Roche). Libraries were checked for quality and concentration using the DNA High-Sensitivity TapeStation assay (Agilent Technologies) and quantitative PCR (Roche), according to the manufacturers’ instructions. Libraries were sequenced 100 bp paired-end on an Illumina NovaSeq 6000 using the S2 Reagent Kit v1.5.

### RNA sequencing (RNA-seq)

RNA-seq data was generated from mESCs derived from 10 mouse strains (129S1/SvImJ, A/J, C57BL/6J CAST/EiJ, NOD/ShiLtJ, NZO/HILtJ, PWK/PhJ, WSB/EiJ, C3H/HeJ, and DBA/2J). Tissues were lysed and homogenized in TRIzol Reagent (Ambion), then RNA was isolated using the miRNeasy Mini kit (Qiagen), according to manufacturers’ protocols, including the optional DNase digest step. RNA concentration and quality were assessed using the Nanodrop 8000 spectrophotometer (Thermo Scientific) and the RNA ScreenTape Assay (Agilent Technologies). Libraries were constructed using the KAPA mRNA HyperPrep Kit (Roche Sequencing and Life Science), according to the manufacturer’s protocol. Briefly, the protocol entails isolation of polyA containing mRNA using oligo-dT magnetic beads, RNA fragmentation, first and second strand cDNA synthesis, ligation of Illumina-specific adapters containing a unique barcode sequence for each library, and PCR amplification. The quality and concentration of the libraries were assessed using the D5000 ScreenTape (Agilent Technologies) and Qubit dsDNA HS Assay (ThermoFisher), respectively, according to the manufacturers’ instructions. Libraries were sequenced 100 bp paired-end on an Illumina NovaSeq 6000 using the S2 Reagent Kit v1.5.

### Genome assembly, variant calling, and QC

For each sample, raw PacBio CLR bam files were converted to FASTA files with bam2fastx (v1.3.0). Strain-specific assemblies were then constructed with Flye (v2.8.3) and Arrow (v2.0.2) was used for polishing. Assembly metrics were computed and summarized with quast (v5.02) (Table S1).

Variant discovery was performed against the GRCm39 (mm39, *mus musculus*) reference genome. SVs were called with PAV (v1.1.2) using minimap2 (v2.17) alignments. Orthogonal callset support was obtained with pbsv (v2.6.2), Sniffles (v1.0.22), SVIM (v2.0.0), SVIM-asm (v1.0.2), and PAV run with the LRA (v1.3.0) aligner (PAVLRA). For pbsv and SVIM, we utilized the pbmm2 (v1.4.0) alignment tool to align raw long reads to the GRCm39 (mm39, *mus musculus*) genome. For Sniffles, NGMLR (v0.2.7) alignments were used. Additional raw-read support was annotated by extracting aligned segments of long-reads in regions around SVs with subseq (Ebert *et al*., 2021) (2+ supporting reads).

For short-read SV calling, raw Illumina FASTQ files were aligned to the GRCm39 reference genome with BWAMEM (v0.7.17). We then called SVs with Manta (v1.6.0), LUMPY (v0.2.13), and DELLY (v0.8.7).

Variants were accepted into the final callset if they have a) either pbsv or SVIM support in addition to raw-read subseq support, b) support from two or more of Sniffles, SVIM-asm, and PAV-LRA in addition to raw-read subseq support, c) both SVIM-asm and PAV-LRA support and are greater than 250 bp, or d) They have raw-read subseq support or duphold-CLR support, have duphold-SR support, and have support from one or more of DELLY, LUMPY, or Manta. The full reference assembly was present during read and contig alignments, however, the final callset was filtered to only include chromosome scaffolds excluding chrY (chr1-19,X).

Finally, A nonredundant callset across samples was achieved by merging with SV-Pop (v2.0.0).

### Segmental duplication annotation and callable regions

Segmental duplications (SD) were characterized in GRCm39 using the whole-genome assembly comparison (WGAC) method as previously described (Bailey et al, 2001). Briefly, WGAC identifies non-homologous pairwise alignments by performing an all-by-all comparison after removing common repeat elements (RepeatMasker v4.1.0 with engine crossmatch v1.090518 and the ‘Mus musculus’ and Tandem Repeat Masker v4.09 while ignoring repeats with > 12 periodicity available from the UCSC browser). WGAC fragments the genome into 400 kbp repeat-masked segments and performs an all-by-all comparison between segments (blastall v2.2.11) and within 400 kbp alignments using the LASTZ tool (v 1.02.00) (https://www.bx.psu.edu/~rsharris/lastz/). Common repeats are reintroduced to construct optimal global alignments that are at least > 1 kbp and > 90% sequence identical to define the SD set for the mouse genome.

### Intersection with mouse genomes project callset and subseq validation

Variants were intersected with the Mouse Genomes Project SV release v5 (Project URL: http://www.sanger.ac.uk/resources/mouse/genomes/; FTP: ftp://ftp-mouse.sanger.ac.uk/; and lifted over to GRCm39 coordinates using UCSC LiftOver). Because callset characteristics were different than modern callsets, we adopted a customized approach. Because insertions lengths are unknown, we counted variants as supported if the insertion site was within 250 bp and ignored the size. Large insertions are called as duplicated reference loci, and to allow direct comparisons, we re-mapped insertion sequences to the reference and intersected the resulting loci with duplication calls using 50% reciprocal overlap. Deletions were also matched by 50% reciprocal overlap.

To validate calls with long-read support, we utilized subseq to extract read lengths (200 bp surrounding region of SV) from pbmm2 alignments. Calls were accepted as PASS if the mean read length difference was ≥ 50 bp.

### Transposable element annotation

In order to characterize transposable element variation, we obtained a FASTA output from SV-Pop containing the full sequences of insertion and deletion SV. We then utilized RepeatMasker (v4.1.2) to annotate repeat sequences within these variant sequences. Transposable element variants (TEVs) are accepted as insertion or deletion variants which have 60% of their content match a specific transposable element family. Full length long terminal repeat (LTR) transposable element consensus sequences are fragmented in transposable element databases as their internal sequence is separate from their flanking LTRs. We further annotated LTR TEVs by accepting the flanking LTR annotation if it were adjacent to an internal sequence. TEV annotations are present in the final data table (Table S2), with annotation for both TE type and TE subtype.

### Variant effects and association analysis

A VCF file containing our primary SV callset was used as input for the Ensembl Variant Effect predictor tool (command-line) to intersect SV coordinates with GRCm39 features. Frameshift_variant, inframe_insertion, inframe_deletion, frameshift_variant, stop_gained, stop_lost, protein_altering_variant, stop_retained_variant, and start_lost annotations were then merged into one “coding_sequence_variant” annotation. A VEP summary analysis was performed and is reported in the Table S2 (VEP_SUMMARY column). We performed an association analysis for each variant consequence (https://useast.ensembl.org/info/genome/variation/prediction/predicted_data.html) and four criteria: Tandem Repeat SVs, Transposable element variants, variants which have short-read support, and variants which have support from the Mouse Genomes Project. For each criterion, we performed a Fisher’s exact test to determine contingency of each SV consequence and criteria (Table S4).

### Chromatin accessibility, gene expression profiling, and transposable element association

Open chromatin peaks were detected for each sample by using the nextflow atacseq (nf-core/atacseq v1.2.1) pipeline. Raw read counts for each open chromatin region were extracted by using featureCounts (v2.0.1). For each sample, regions surrounding SV coordinates (± 2500 bp flanking the breakpoint) were intersected with open chromatin regions using BEDTools intersect (v 2.30.0). The sum of raw read counts for overlapping peaks were accepted for further analysis. Read count values for each sample was merged into a matrix and normalized with DESeq2 (v3.15). For each SV, normalized read counts for each sample were grouped into two groups dependent on if the sample contained the SV or not and a Mann-Whitney U test was performed between each group to determine a statistically significant *(p* < 0.05) difference. To assess TE families and changes in chromatin accessibility association, SVs which were associated with changes in chromatin accessibility were first grouped by TE subtype. Subtype-specific calls were further grouped whether they were an insertion or deletion variant, and a Student’s *t*-test was performed between normalized read count values for insertion and deletions of a particular TE subtype, followed by Bonferroni correction.

For RNA-seq analysis, raw FASTQ files were aligned to GRCm39 reference with STAR (v2.7.8a). Raw gene read counts were obtained for each sample with the TEtranscripts package (v2.2.1) and normalized with the DESeq2 package (v3.15). Association between candidate SVs and gene expression changes were calculated by a nonparametric test (Mann-Whitney U) of normalized gene read counts between samples which contained the candidate SV and those that did not. For transcript assemblies, we utilized the StringTie package (v2.1.7).

### PCR validations and false discovery rate

For each SV, 500 bp of flanking DNA sequence was obtained from selected SV breakpoints using BEDTools getfasta. Primers were designed using Primer3web (version 4.1.0; https://primer3.ut.ee/). Ideal primer conditions were set to have a length of 25 bp, melting temperature of 65°C, and 50% GC content. Primers were designed to anneal at least 50 bp from SV breakpoints. At least one primer was required to anneal in a unique region of the genome using UCSC BLAT (https://genome.ucsc.edu/cgi-bin/hgBlat), and primer pairs were analyzed using UCSC In-Silico PCR (https://genome.ucsc.edu/cgi-bin/hgPcr) to confirm a unique amplicon for each SV. If primer conditions were not met in the first 500 bp flanking region, the design window was increased in increments of 500 bp until all primer requirements are fulfilled. Takara LA Taq Polymerase (Takara RR002M) was used for PCR validation. Each primer pair was tested across 8 founder mice strains and a negative control. Touchdown PCR was used for desirable primer annealing and amplification of each predicted amplicon. OrgangeG gel loading dye was mixed with PCR product and ran in a 1% agarose gel (Bio-Rad Certified Molecular Biology Agarose 1613102) supplemented with ethidium bromide (Bio-Rad 1610433). Each primer pair was designed to show a resulting amplicon whether or not the SV was present. PCR validations and primer sequences can be found in Table S3.

In order to provide an unbiased FDR estimate, variant callset filters were informed by inspecting callset support from all sources and identifying sources of likely false SV calls, was not informed by PCR validation results, and adjustments to the filters were not made once FDR estimates were computed.

Thermocycler program for amplicon sizes less than 5 kbp:

Step 1. 95°C for 1 min

Step 2. 95°C for 30 sec

Step 3. 68°C for 30 sec (with a 1°C ramp down per cycle) Step 4. 72°C for 5 min

Step 5. Return to step 2 and repeat for 5 cycles Step 6. 95°C for 30 sec

Step 7. 63°C for 30 sec

Step 8. 72°C for 5 min

Step 9. Return to step 6 and repeat for 25 cycles Step 10. 72°C for 10 min

Step 11. 4°C infinite hold

Thermocycler program for amplicon sizes greater than 5 kbp:

Step 1. 95°C for 1 min

Step 2. 95°C for 30 sec

Step 3. 68°C for 30 sec (with a 1°C ramp down per cycle) Step 4. 72°C for 9 min

Step 5. Return to step 2 and repeat for 5 cycles Step 6. 95°C for 30 sec

Step 7. 63°C for 30 sec

Step 8. 72°C for 9 min

Step 9. Return to step 6 and repeat for 25 cycles Step 10. 72°C for 10 min

Step 11. 4°C infinite hold

## Acknowledgements

This research was supported by The Jackson Laboratory Director’s Innovation Fund, The Jackson Laboratory Cancer Center (P30CA034196), the National Institutes of Health (R35 GM133600 to C.R.B., R24 OD021325-06 to L.G.R., R01 HG002385 to E.E.E., and 1T32HG010463 to AF). We would like to thank Charles Lee for his review of the manuscript and Tonia Brown for her critical proofreading assistance. We would also like to thank The Jackson Laboratory Genome Technologies core for help with sequencing. E.E.E. is an investigator of the Howard Hughes Medical Institute.

